# Genotyping and epidemiological metadata provides new insights into population structure of *Xanthomonas* isolated from walnut trees

**DOI:** 10.1101/397703

**Authors:** Camila Fernandes, Pedro Albuquerque, Leonor Cruz, Fernando Tavares

**Author notes:** Address correspondence to Camila Fernandes, or Fernando Tavares,. Present address: Pedro Albuquerque, I3S - Instituto de Investigação e Inovação em Saúde, Universidade do Porto, Porto, Portugal.

## Abstract

*Xanthomonas arboricola* pv. *juglandis* (*Xaj*) is the etiological agent of walnut diseases affecting leaves, fruits, branches and trunks. Although this phytopathogen is widely spread in walnut producing regions and has a considerable genetic diversity, there is still a poor understanding of its epidemic behaviour. To shed some light on the epidemiology of these bacteria, 131 *Xanthomonas* isolates obtained from 64 walnut trees were included in this study considering epidemiological metadata such as year of isolation, bioclimatic regions, walnut cultivars, production regimes, host walnut specimen and plant organs. Genetic diversity was assessed by multilocus sequence analysis (MLSA) and dot blot hybridization patterns obtained with nine *Xaj*-specific DNA markers (XAJ1 – XAJ9). The results showed that *Xanthomonas* isolates grouped in ten distinct MLSA clusters and in 18 hybridization patterns (HP). The majority of isolates (112 out of 131) were closely related with *X. arboricola* strains of pathovar *juglandis* as revealed by MLSA (clusters I to VI) and hybridize with more than five *Xaj*-specific markers. Nineteen isolates clustered in four MLSA groups (clusters VII to X) which do not include *Xaj* strains, and hybridize to less than five markers. Taking this data together, was possible to distinguish 17 lineages of *Xaj*, three lineages of *X. arboricola* and 11 lineages of *Xanthomonas* sp. Some *Xaj* lineages appeared to be widely distributed and prevalent across the different bioclimatic regions and apparently not constrained by the other features considered. Assessment of type III effector genes and pathogenicity tests revealed that representative lineages of MLSA clusters VII to X were nonpathogenic on walnut, with exception for strain CPBF 424, making this bacterium particularly appealing to address *Xanthomonas* pathoadaptations to walnut.

**IMPORTANCE:** *Xanthomonas arboricola* pv. *juglandis* is one of the most serious threats of walnut trees. New disease epidemics caused by this phytopathogen has been a big concern causing high economic losses on walnut production worldwide. Using a comprehensive sampling methodology to disclose the diversity of walnut infective *Xanthomonas*, we were able to identify a genetic diversity higher than previously reported and generally independent of bioclimatic regions and the other epidemiological features studied. Furthermore, co-colonization of the same plant sample by distinct *Xanthomonas* strains were frequent and suggested a sympatric lifestyle. The extensive sampling carried out resulted in a set of non-*arboricola Xanthomonas* sp. strains, including a pathogenic strain, therefore diverging from the nonpathogenic phenotype that have been associated to these atypical strains, generally considered to be commensal. This new strain might be particularly informative to elucidate novel pathogenicity traits and unveil pathogenesis evolution within walnut infective xanthomonads. Beyond extending the present knowledge about walnut infective xanthomonads, this study might contribute to provide a methodological framework for phytopathogen epidemiological studies, still largely disregarded.

## Introduction

*Xanthomonas* is a gammaproteobacteria genus belonging to Xanthomonaceae, which includes soil dwelling bacteria and important phytopathogenic species, often composed of several host-specific pathovars (1, 2). *X. arboricola* pv. *juglandis* (*Xaj*) is the only xanthomonads described as a pathogen on walnut trees, acknowledged as the etiological agent of walnut pathologies known as the “Walnut Bacterial Blight (WBB); the “Brown Apical Necrosis” (BAN); and the “Vertical Oozing Canker” (VOC), all causing considerable economic losses (3-6). *Xaj* is frequently isolated from symptomatic walnut leaves and fruits, but also from other walnut plant organs without apparent visible lesions, namely dormant walnut buds and catkins (7, 8), and from non-plant materials, such as orchard machinery (9). This pathogen has a cosmopolitan distribution and diseased trees are commonly observed throughout walnut-growing regions (10, 11). Genotyping studies have found a considerable genetic diversity in *Xaj*, which is reportedly high when compared with the diversity described for other *X. arboricola* pathovars (5, 11-14). This feature, together with evidence of genomic trade-offs within the species (5, 13, 15, 16), evokes an opportunistic pathogen, even though evidence for environmental reservoirs of *Xaj* remains poorly characterized (11, 17-19). These data are further supported by the isolation of nonpathogenic strains of *X. arboricola*, described as phylogenetically heterogeneous, and grouping separately from the well-defined clusters of pathogenic *Xaj* strains (20). Assessment of *Xaj* in France (5), Serbia (21) and Italy (9) showed large genetic diversity among isolates and proposed the existence of diverse *<u>Xaj</u>* populations. Furthermore, the genetic differences found between *Xaj* strains collected from VOC and WBB symptoms, suggests the presence of distinct genetic lineages within *Xaj* populations (5).

Different factors have been hypothesized to explain *Xaj* population diversity, namely geographical location (12, 22); origin of plant propagation material (5, 9); adaptation to particular environmental conditions (22, 23); genome flexibility or pathogen virulence (21-23); or even selective pressure by the host plant (5, 24). Regardless their valuable contributions, these studies were either based on a low number of bacterial isolates, often obtained without a planned sampling strategy, or based on a set of *Xaj* strains from worldwide collections, overlooking important metadata such as date, plant host traits, and climatic features which are essential to determine epidemiological patterns (25, 26). In fact, to understand the epidemiological behaviour of *Xaj*, it is of utmost importance to conciliate in a single study comprehensive genotyping analysis of a coherent set of isolates with insightful metadata.

This study aimed to characterize the genetic diversity of xanthomonads isolates obtained from walnut trees over a three-year period, across distinct bioclimatic regions and taking into account different walnut cultivars, production regimes (orchards vs. isolated trees), host walnut tree and plant organs. Multilocus sequence analysis (MLSA) and dot blot hybridization patterns of nine *Xaj*-specific markers (27), were used to determine the genetic diversity of isolates. These data were complemented by monitoring the presence/absence of four informative type III effector genes (T3E) observed to differentiate non-pathogenic and pathogenic *Xaj* clusters (20) and by pathogenicity tests of representative lineages. Altogether, this research contributes to better elucidate the impact of environmental factors and host features on bacterial population diversity, which is important to improve phytosanitary control of diseases caused by walnut pathogenic *Xanthomonas*. Furthermore, we believe that the current study provides a methodological framework to address epidemiological studies of phytopathogens.

## Results

### Bacterial isolates obtained from walnut trees

A total of 131 isolates displaying yellow-pigmented colonies typical of *Xanthomonas* species were obtained. The majority of isolates were collected from symptomatic plant material (94%, 122/131), mostly from leaf samples (97 isolates, 74%), whereas nine (6%, 9/131) isolates were obtained from asymptomatic plant organs (Table 1). More than one isolate was recovered from 38 trees, either from the same sampling occurrence (i.e. the same plant in the same date) or from the same walnut tree at different sampling dates (Table 1).

**TABLE 1.**
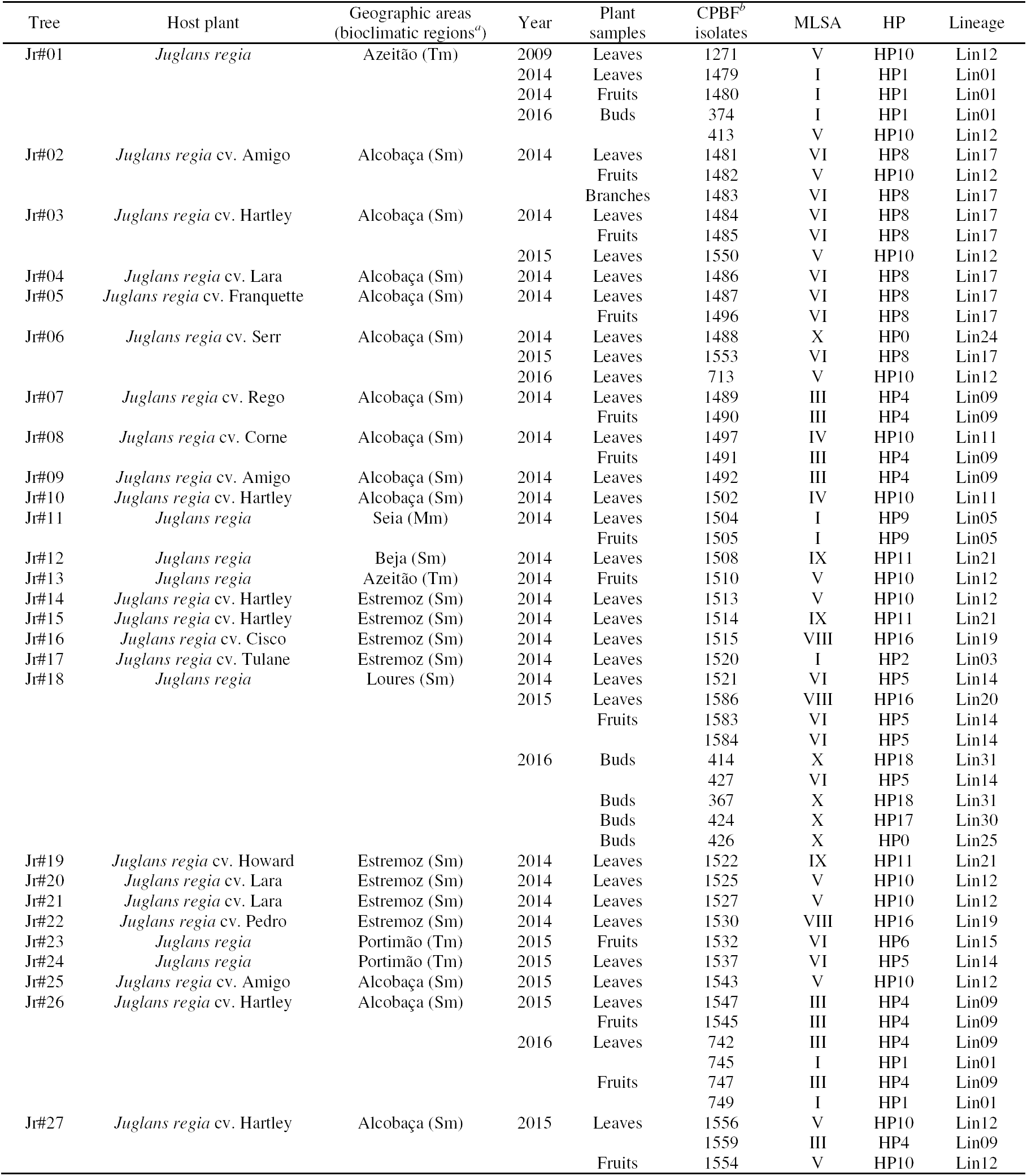
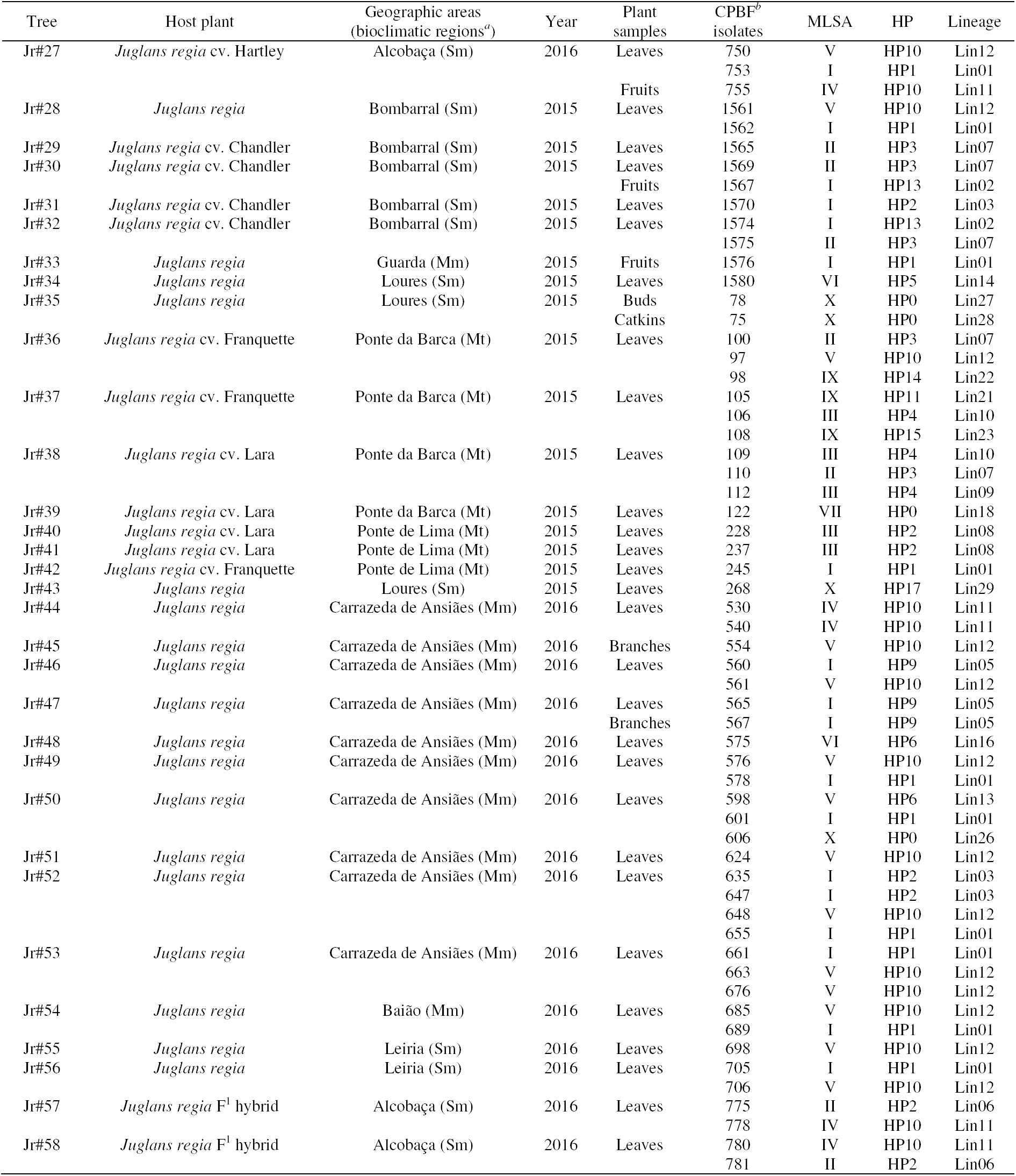
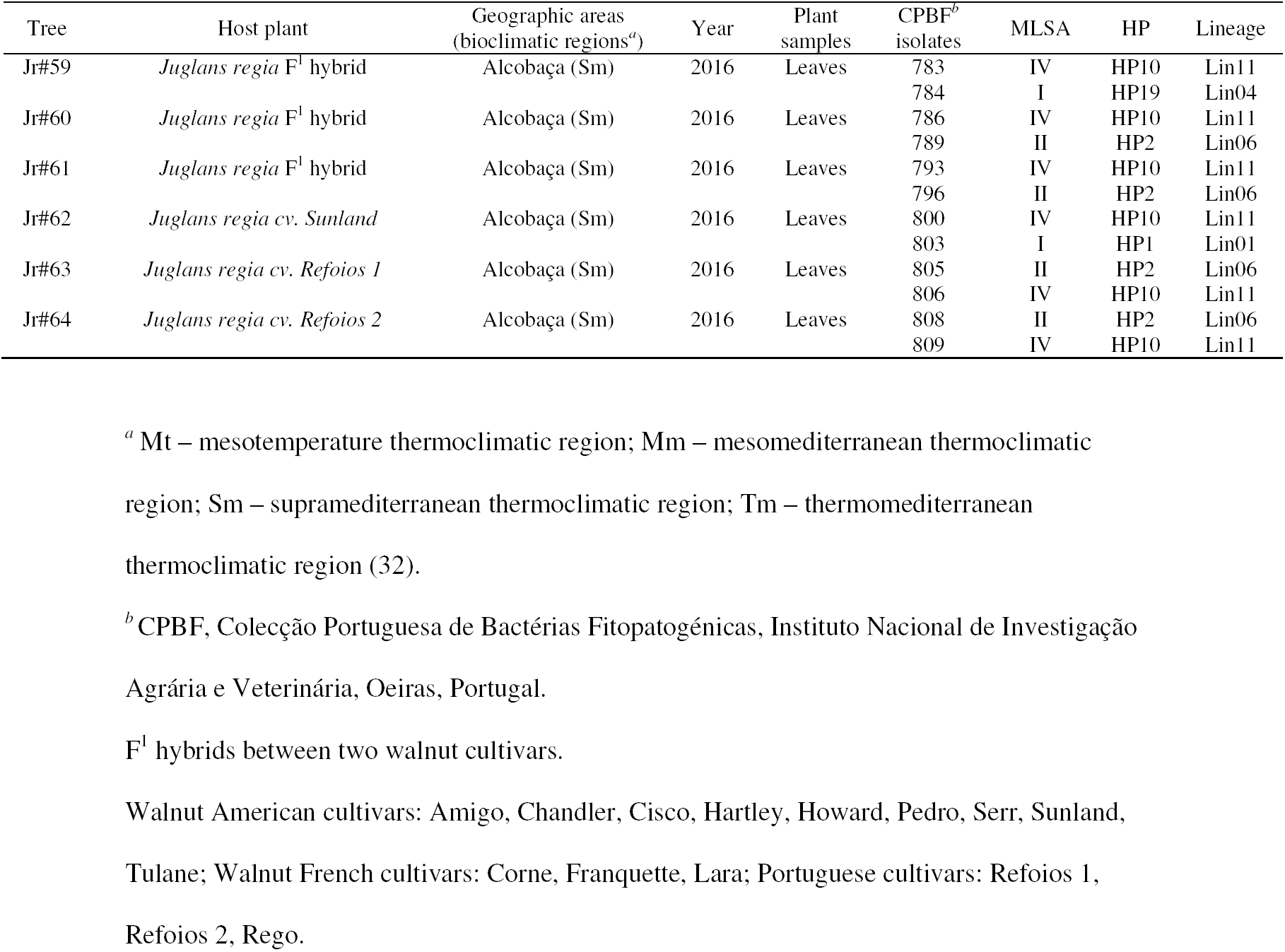
Walnut trees included in this study and the correspondent epidemiological information and molecular results obtained.

### MLSA revealed that *Xanthomonas* walnut isolates were grouped into ten distinct clusters

The MLSA clustering data showed a clear separation, supported by high bootstrap values of 99%, between the 131 isolates and strains representatives of different *Xanthomonas* species, namely *X. albilineans, X. alfalfae, X. axonopodis, X. sacchari, X. campestris, X. cassava*e, *X. citri, X. euvesicatoria, X. floridensis, X. fragariae, X. fuscans, X. gardneri, X. hortorum, X. oryzae, X. perforans, X. prunicola, X. translucens, X. vasicola*. Even so, all CPBF isolates were assigned into the genus *Xanthomonas* and were clustered in ten different groups (Clusters I to X, Fig. 1). Most isolates (85.5%, 112/131) were grouped in clusters I to VI together with strains of *X. arboricola* pv. *juglandis* and other closely related strains belonging to pathovars *pruni* and *corylina*. The other 19 (14,5%, 19/131) isolates belong to clusters VII to X. Isolates from clusters VII and VIII were closely related with *X. arboricola* CFBP 7634 and *X. arboricola* CFBP 7651, previously described as nonpathogenic strains by Essakhi et al. (20); with *X. arboricola* CITA 124, *X. arboricola* CITA 44 and *X. arboricola* CITA 14, previously described as avirulent strains by Garita-Cambronero et al. (28, 29) and with strain NCPPB 1630 belonging to the poorly characterized pathovar *celebensis* (30) and *X. arboricola* 3004 isolated from barley and with uncertain pathogenicity (31). Isolates of groups IX and X formed two different clusters (bootstrap values > 95%) distant from the other isolates and strains of *X. arboricola* (Fig. 1). Cluster IX is composed of six isolates (CPBF 98, CPBF 105, CPBF 108, CPBF 1508, CPBF 1514, CPBF 1522) showing identical MLSA concatenated sequences, whereas considerable nucleotide differences were observed among seven (CPBF 75, CPBF 78, CPBF 268, CPBF 424, CPBF 426, CPBF 606, CPBF 1488) of the nine isolates grouped into cluster X.

**FIG 1.**
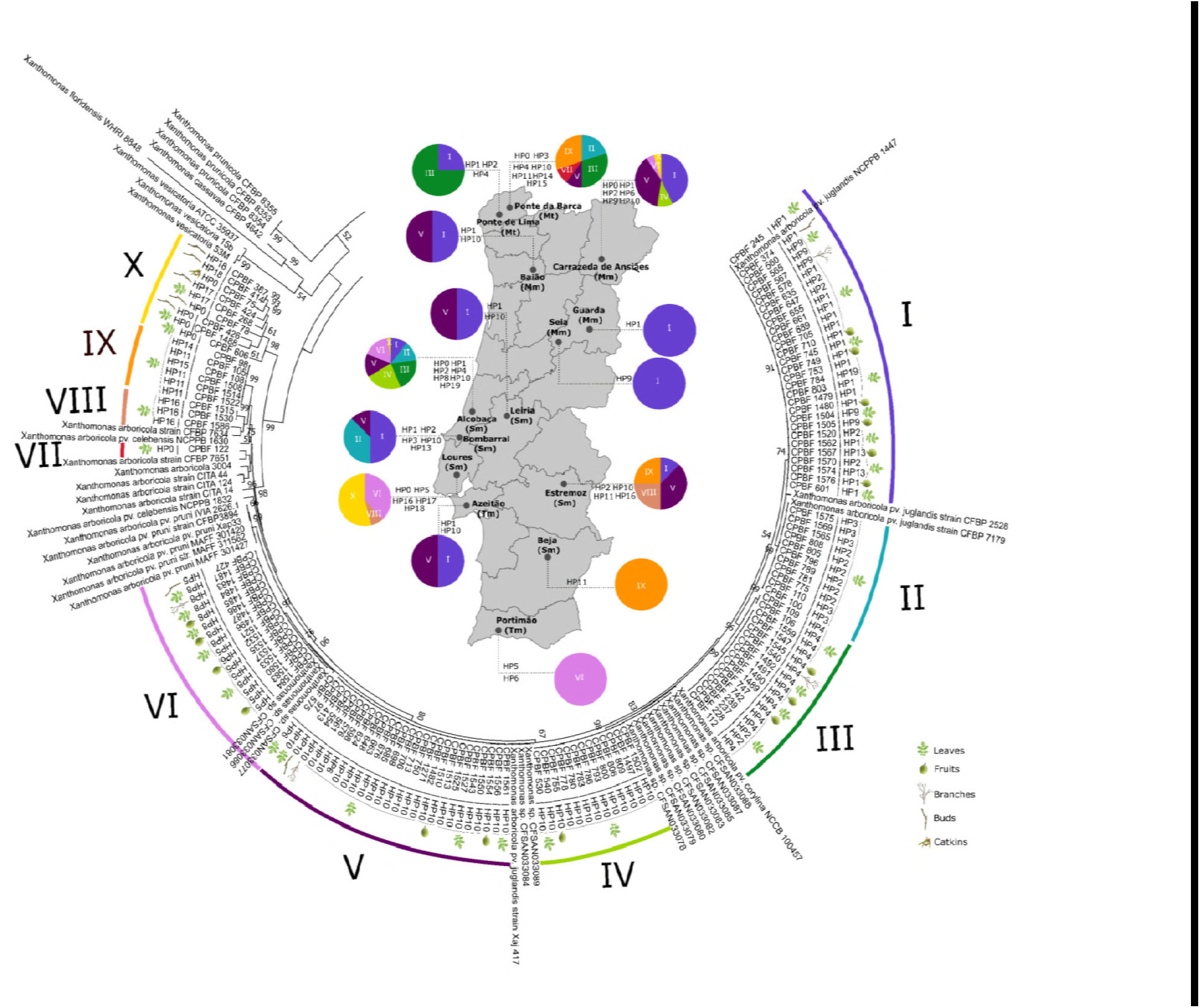
MLSA analysis and dot blot genotyping coupled with metadata including geographic locations, bioclimatic regions, and walnut host organs for the 131 *Xanthomonas* walnut isolates used in this work. The Maximum Likelihood tree was based on the nucleotide alignments of 252 concatenated sequences (2774 bp) of *acnB, gyrB, fyuA* and *rpoD* genes, using the General Time Reversible (GTR+G+I) model. Bootstrap values higher than 50 are shown. The tree was edited using MEGA 7.0 (40) and the principal results are showed. Distinct MLSA clusters (I to X) of *Xanthomonas* isolates are highlighted with different colours. For all the isolates, the respective hybridization patterns (HP0 to HP19) is shown, as well as the plant organ from which they were isolated. The map in the centre highlights the fourteen geographic locations sampled, and details the respective thermoclimatic classification: Mt – mesotemperature; Mm – mesomediterranean; Sm – supramediterranean; Tm – thermomediterranean (32). The coloured pie charts indicate the prevalence of MLSA clusters and identifies the different hybridization patterns (HP) for each sampled region.

### Diversity of isolates assessed using *Xaj* specific DNA markers

A total of 18 different hybridization patterns (HP), identified for the nine *Xaj* specific markers (XAJ1 to XAJ9) were obtained for the 131 isolates (Fig. 1 and Fig. 2). The most representative HP was HP10, identified in 38 isolates (29%, 38/131). The less frequent HPs were HP14, HP15 and HP19, only identified in one isolate each, followed by HP6, HP11, HP16, HP17, HP18 found in less than five isolates (Fig. 1). Among these, HP16 (CPBF 1515, CPBF 1586 and CPBF 1530), HP17 (CPBF 268 and CPBF 424) and HP18 (CPBF 414 and CPBF 367) correspond to hybridization patterns limited to a single marker, XAJ9, XAJ2 and XAJ1, respectively. Six isolates out of 131 isolates (CPBF 75, CPBF 78, CPBF 122, CPBF 426, CPBF 606 and CPBF 1488) provided negative hybridization results for all markers (pattern HP0, Fig. 1 and Fig. 2).

**FIG 2.**
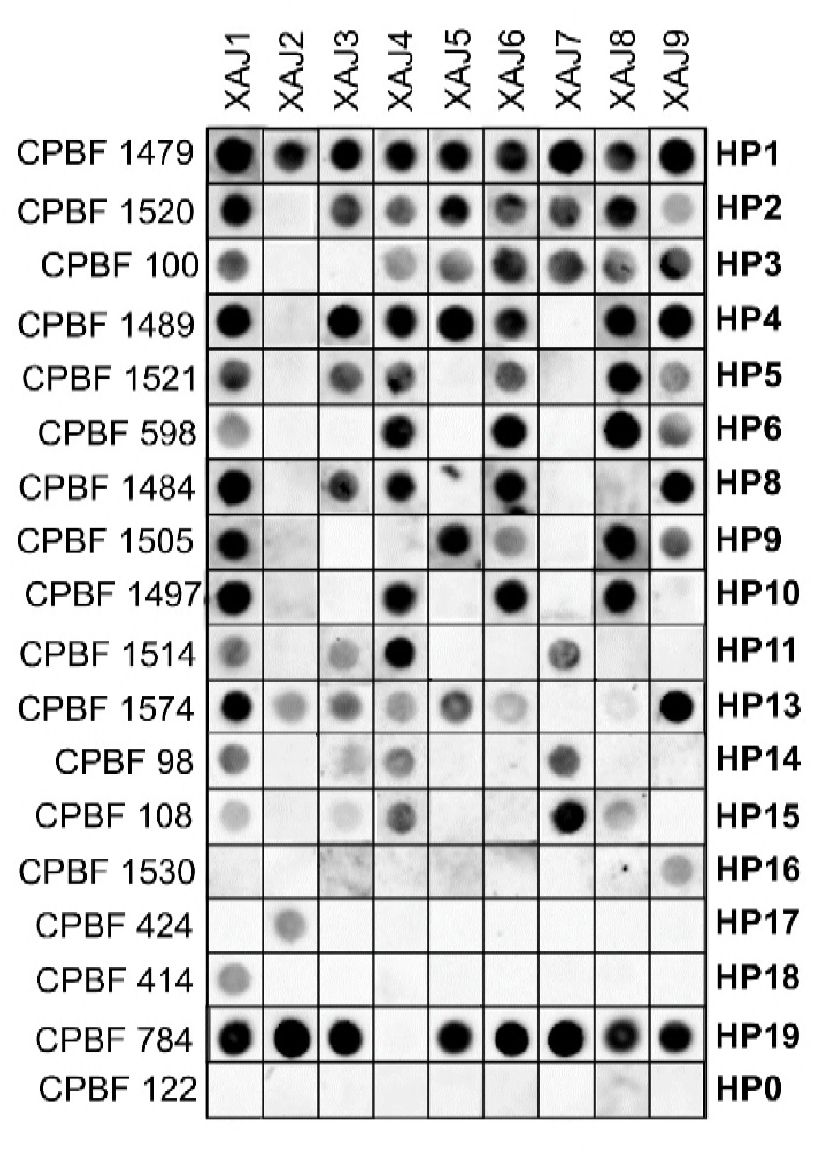
Dot blot matrix summarizing the 18 different hybridization patterns (HP0 to HP19) obtained with the 131 isolates using nine *Xanthomonas arboricola* pv. *juglandis* specific markers (XAJ1 to XAJ9). Strain LMG 751 was used as positive control for all dot blot assays.

### Dot blot and MLSA data allowed to distinguish 31 xanthomonads lineages

Patterns characterized by hybridization to three or less markers (HP0, HP14, HP16, HP17 and HP18, Fig. 2) were exclusive to strains of MLSA clusters VII to X (Fig. 1). On the other hand, strains belonging to MLSA groups I, II and III hybridized to seven or more markers (HP1, HP2, HP3, HP4, HP13, HP19, Fig. 1 and Fig. 2), except for five strains belonging to MLSA group I which hybridize to five markers (HP9, Fig. 1 and Fig. 2).

For the 131 isolates obtained in this work, a total of 20 unique concatenated sequences were obtained with 403 polymorphisms in 199 nucleotide sites, taking into account MLSA data, and 18 different hybridization patterns were obtained by dot blot assays. The added discrimination of the dot blot analysis, based on the presence/absence of specific loci, allowed the following group subdivisions: Group I with five distinct clonal lineages, Group II with two lineages, Group III with three lineages, Group IV with no subdivisions (one lineage), Group V with two lineages, Group VI with four lineages, Group VIII with two lineages, Group IX with three lineages and Group X with eight lineages. Group VII comprised only one strain (one lineage). In total, the compilation of data from the two genotyping strategies allowed distinguishing 31 lineages within the xanthomonads populations found in walnut trees (Table 1). Lineages 1 to 17 were represented by *Xaj* isolates, lineages 18 to 20 were represented by *X. arboricola* isolates and lineages 21 to 31 were represented by *Xanthomonas* sp. isolates.

Eight distinct genetic lineages (Lin1, Lin3, Lin7, Lin9, Lin11, Lin12, Lin14 and Lin21) were found across the 14 geographic locations of Portugal analysed in this study. The same lineage was found in locations characterized by distinct thermoclimatic regions according Rivas-Martínez (32): e.g. Lin1, Cluster I/HP1, was found in Ponte de Lima, a mesotemperature region; in Carrazeda de Ansiães, a mesomediterranean region; in Alcobaça, a supramediterranean region; and in Azeitão, a thermomediterranean region. Twenty lineages (Lin2, Lin4, Lin8, Lin10, Lin13, Lin15, Lin16, Lin18, Lin19, Lin20, Lin22, Lin23, Lin24, Lin25, Lin26, Lin27, Lin28, Lin29, Lin30, and Lin31) were represented by one or two isolates. Additionally, isolates belong to lineages Lin6 (cluster II/HP2) and Lin17 (cluster VI/HP8) were restricted to one specific geographic location (Alcobaça) and Lin5 (cluster I/HP9) was only found in the mesomediterranean interior region of Portugal (Seia and Carrazeda de Ansiães locations). Moreover, five lineages were always found in geographic locations where walnut trees were sampled in consecutive years (Lin9 and Lin12 in Alcobaça, Lin1 and Lin12 in Azeitão and Lin14 in Loures). None of the most representative lineages were associated with specific plant organs, except for lineage Lin6 represented by six isolates only found in leaves, or with specific *J. regia* cultivars.

### Isolates with distinct genotypes colonize the same organ of walnut trees

From the 64 trees analysed in this work, more than one isolate was retrieved from 37 trees sampled at the same date (Table 1, Jr#01, Jr#02, Jr#03, Jr#05, Jr#07, Jr#08, Jr#11, Jr#18, Jr#23, Jr#25, Jr#26, Jr#27, Jr#28, Jr#30, Jr#32, Jr#35, Jr#36, Jr#37, Jr#38, Jr#41, Jr#44, Jr#46, Jr#47, Jr#49, Jr#50, Jr#52, Jr#53, Jr#54, Jr#56, Jr#57, Jr#58, Jr#59, Jr#60, Jr#61, Jr#62, Jr#63 and Jr#64). From these, five trees allowed to recover only one genetic lineage, as defined using MLSA and HP data (Jr#05, cluster VI/HP8, Lin17; Jr#07, I/HP4, Lin9; Jr#11, I/HP9, Lin5; Jr#44, IV/HP10, Lin11; Jr#47, I/HP9, Lin5), while 32 trees allowed to discern two or more distinct bacterial lineages from a single sampling event, with the majority of isolates being obtained from the same leaf samples of walnut trees (22 trees: Jr#26, Jr#27, Jr#28, Jr#32, Jr#36, Jr#37, Jr#38, Jr#46, Jr#49, Jr#50, Jr#52, Jr#53, Jr#54, Jr#56, Jr#57, Jr#58, Jr#59, Jr#60, Jr#61, Jr#62, Jr#63, Jr#64, Fig. 3). Interestingly, all six trees that were analysed in consecutive years (Jr#01, Jr#03, Jr#06, Jr#18, Jr#26, Jr#27), consistently allowed recovering different lineages (Table 1 and Fig. 1).

**FIG 3.**
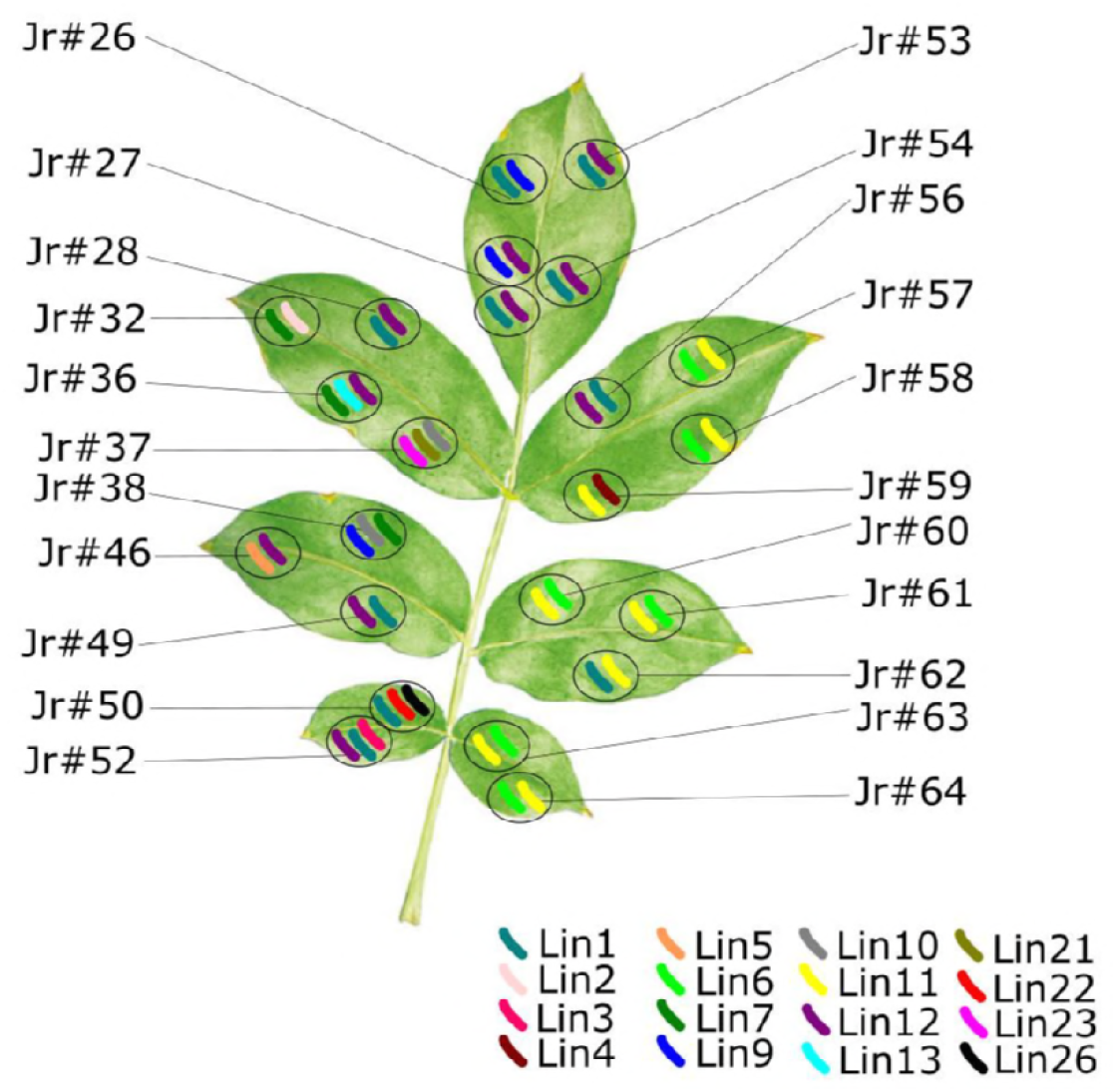
Schematic representation of co-colonization (highlighted by circles) by different xanthomonads lineages assigned to different colours on leaves of 22 walnut specimens (Jr#) during the sampling period (2014 – 2016). Lin1 to Lin13 correspond to *Xaj* isolates. Lin21 to Lin26 correspond to non-*arboricola Xanthomonas* sp. isolates.

### Assessment of type III efector genes (*xopR, avrbs2, xopF1* and *xopN*), revealed *Xanthomonas* isolates with new T3E profiles

The presence of T3E genes (*xopR, avrbs2, xopF1* and *xopN*) was assessed on representative isolates of each main cluster constituted by *Xaj* isolates (clusters I to VI) and on all *X. arboricola* isolates belonging to clusters VII and VIII and *Xanthomonas* sp. from cluster IX and X (Fig. 1 and Fig. S1). Among CPBF *Xaj* isolates belonging to clusters I to VI, positive hybridization dots were obtained for all four T3E genes studied. *xopN* was not detected in all the 19 isolates belonging to MLSA clusters VII, VIII, IX and X. Ten of these isolates, belonged to clusters VII, VII and IX, and hybridized with *xopR, avrBs2* and *xopF1* specific probes. Two isolates of cluster X (CPBF 78 and CPBF 424) hybridized with *xopR* and *xopF1* and the remaining seven isolates of cluster X only hybridized with *xopR* gene (Fig. 4).

**FIG 4.**
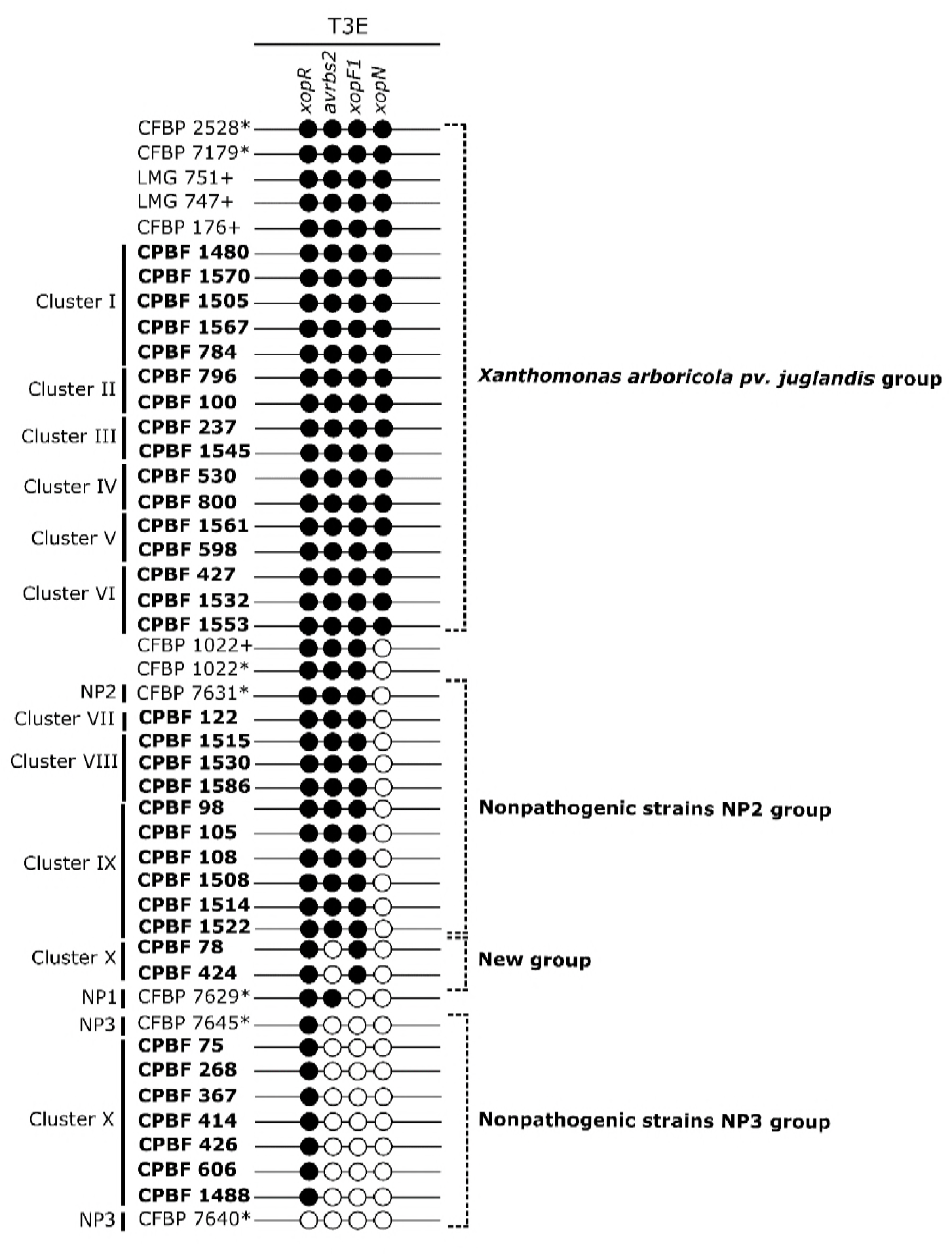
Dot blot matrix reporting the presence/absence of four type three effector genes (T3E - *xopR, avrBs2, xopF1* and *xopN*) in *Xanthomonas* strains and isolates representing the ten MLSA clusters (I to X) and the 18 hybridization patterns (HP0 to HP19). Black circles represent positive dot blot hybridization to the target gene and white circles represent negative dot blot hybridization to the target gene. Sixteen isolates belonging to MLSA clusters I to VI and all isolates grouped in MLSA clusters VII to X were screened. *Xanthomonas arboricola* pv. *juglandis* (*Xaj*) reference strains LMG 751, LMG 747, CFBP 176 and *Xanthomonas arboricola* CFBP 1022 were used as controls (highlighted with +). Dot blot images are supplied in Fig. S1 in the supplemental material. For comparison, the distribution of *xopR, avrBs2, xopF1* and *xopN* genes, described by Essakhi et al. (20) for six strains is highlighted with an asterisk (*), including two *Xaj* strains (CFBP 2528 and CFBP 7179) and four strains isolated from walnut and belonging to three nonpathogenic groups (NP1 – strains CFBP 1022, CFBP 7629; NP2 – strain CFBP 7631; and NP3 – strain CFBP 7645).

The T3E gene patterns obtained for the 35 isolates assayed could be correlated with data previously reported by Essakhi et al. (20), with exception of two isolates (CPBF 78 and CPBF 424), which presented a novel profile of T3E genes (Fig. 4).

### Non-*arboricola Xanthomonas* isolate can cause bacterial leaf spots on walnut

Pathogenicity assays were conducted in order to verify the ability of *Xanthomonas* sp. isolates with distinct T3Es (CPBF 1514, CPBF 424, CPBF 75, CPBF 367 and CPBF 1488) to cause typical disease symptoms on *Juglans regia*. Seven days after inoculation of strain LMG 747, bacterial necrotic spots were observed on leaves of walnut plantlets. Similar symptoms were detected for *Xaj* isolate CPBF 1480 and for the non-*arboricola Xanthomonas* isolate CPBF 424 (Fig. 5). No symptoms were observed using *Xanthomonas* sp. isolates CPBF 1514, CPBF 75, CPBF 367 and CPBF 1488 (data not shown), suggesting they are nonpathogenic on walnut trees. Re-isolation of yellow mucoid bacterial colonies followed by sequencing analysis of two genes (*gyrB* and *fyuA* partial sequences) confirmed that the two isolates (CPBF 1480, a *Xaj* isolate and CPBF 424, *Xanthomonas* sp. isolate) are pathogenic on *Juglans regia* (data not shown).

**FIG 5.**
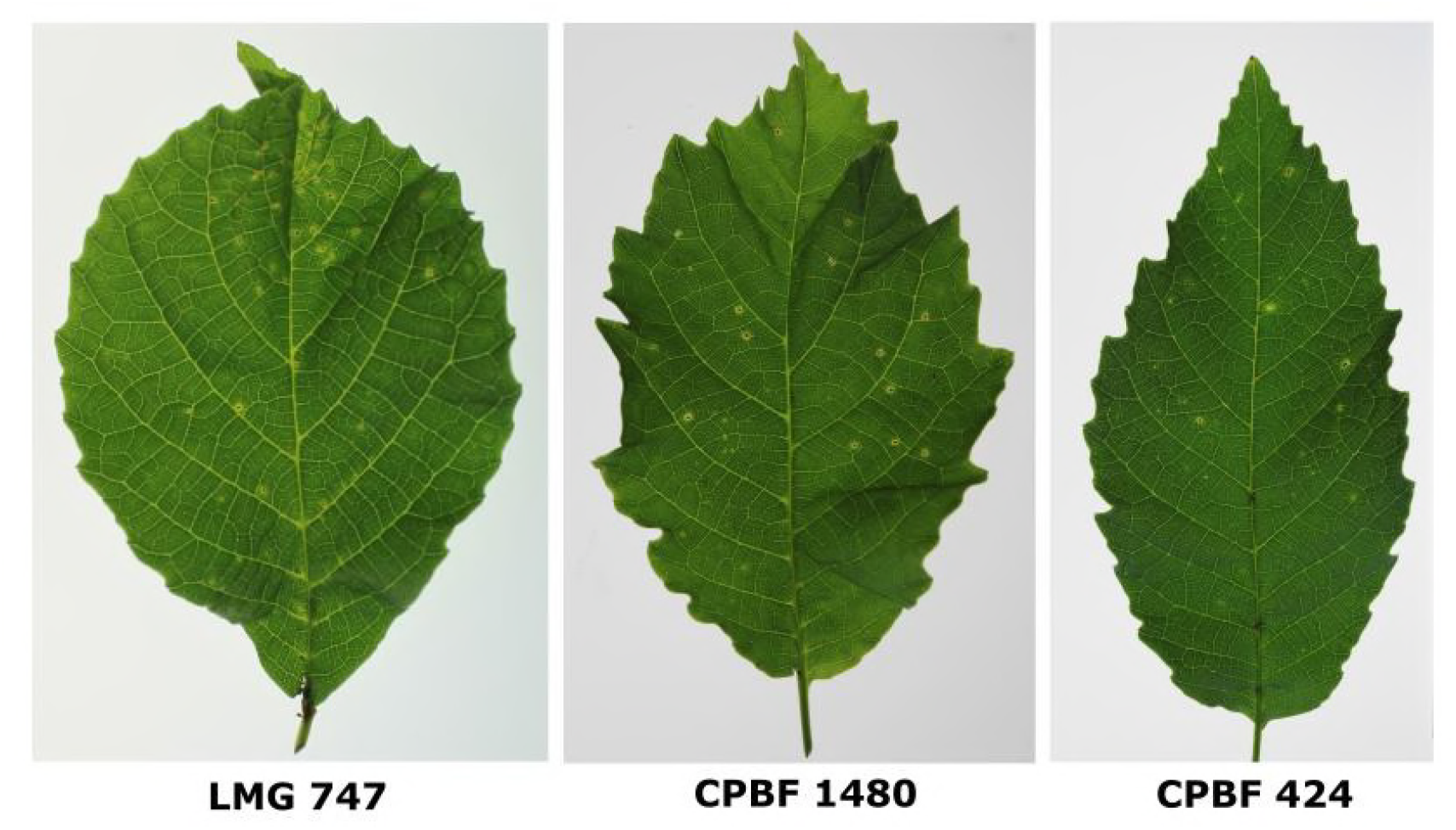
Pathogenicity assays showing symptoms on walnut leaves developed four weeks after inoculation. LMG 747 and CPBF 1480 (MLSA cluster I and HP1) were used as positive controls. Walnut leaves inoculated with strain CPBF 424 (MLSA cluster X) showed similar bacterial necrotic spots as the positive controls. No symptoms were observed for *X. arboricola* isolates CPBF 75, CPBF 367, CPBF 1488 (MLSA cluster X) and for CPBF 1514 (MLSA cluster IX) (data not shown).

## Discussion

Walnut tree is acknowledged as a host plant species for *X. arboricola* providing a permanent niche for the pathovar *juglandis*, which are walnut pathogens characterized by a high genetic diversity (5, 9, 12, 21-24). Regardless these contributions, most of these studies were based either on established collections of strains, or on a limited number of field isolates apparently obtained through a random sampling strategy. To comprehensively characterize the diversity of *Xanthomonas* walnut pathogens, and fully understand the epidemic dynamics of these bacteria, the present research followed a sampling plan taking into account metadata thought to influence disease epidemiology, namely different walnut cultivars, production regimes (orchards vs. isolated trees), walnut tree specimen, plant organs and distinct bioclimatic regions, over a three-year period.

Portugal is a walnut production country characterized by distinct bioclimatic regions and a long historical record of WBB disease symptoms. The first scientific evidence for walnuts infected with *Xaj* date back to 1935 (33). The authors mentioned that high humidity favoured the infection and suggested that both environmental and meteorological conditions could influence disease development (33). Currently, WBB disease is commonly observed in walnut Portuguese orchards and in dispersed walnut trees across the country, affecting crop yield drastically. Although is acknowledged that *Xaj* is the main pathogenic agent of walnut diseases all over the country, information about genetic diversity of this pathogen, its dissemination and epidemiology remain unknown, impairing the implementation of scientifically informed phytosanitary measures. In this work, a total of 131 isolates were obtained from diseased walnut trees between 2014 to 2016 following a planned sampling strategy. Isolates were collected from distinct plant organs (leaves, fruits, branches, buds and catkins) of different cultivars of walnut trees mainly located in geographic regions with high walnut production and characterized by distinct bioclimatic conditions and different management strategies.

Isolates diversity was assessed by MLSA considering four housekeeping genes (*acnB, fyuA, gyrB* and *rpoD*), following an approach previously used to determine the phylogeny of *Xanthomonas* spp. (34, 35). In fact, the large number of sequences available at the GenBank database, the discriminatory power of each of these housekeeping genes and its wide utilization in transversal studies on *Xaj* (9, 21, 23, 24), makes MLSA a suitable method to carry out detailed comparisons with other studies. Furthermore, MLSA has been particularly useful for differentiating *X. arboricola* isolates obtained from the same host plant (20, 28, 29). In this study, ten different MLSA clusters were defined (I to X, Fig.2) with most of the isolates being distributed in clusters I to VI, which also contained several fully sequenced *Xaj* strains, namely *Xaj* 417 (36); NCPPB 1447; CFBP 2528 and CFBP 7179 (37); CFSAN033077-89 (38). The remaining isolates, clustered in groups VII to X (Fig. 2), were clearly distinguishable from the main *Xaj* MLSA clusters. Clusters VII and VIII, closely related to other *X. arboricola*, as non-pathogenic or avirulent strains (20, 28, 29), comprise isolates belong to the species *X. arboricola* isolated from symptomatic leaves. Cluster IX and X were grouped outside any other cluster defined by *X. arboricola*. Cluster IX is formed by a single clonal complex of six *Xanthomonas* sp. isolated from symptomatic leaves and cluster X is the group of *Xanthomonas* sp. mainly isolated from asymptomatic material (bud and catkin samples) showing the highest genetic variation. To further elucidate the genetic diversity of all 131 isolates, dot blot hybridization assays using *Xaj*-specific DNA markers XAJ1 to XAJ9 (27), allowed to identify eighteen distinct hybridization patterns (Fig. 2), and distinguish isolates within the same MLSA cluster (Fig. 1). Particularly, isolates from MLSA cluster I, which included 29 isolates, were divided into four sub-groups when hybridization patterns were considered. The genotyping appraisal of a considerable number of isolates confirm the utility of these markers for the identification of different *Xaj* lineages, as suggested previously (27). Furthermore, these molecular markers have shown to be useful as genetic tools on the characterization of all xanthomonads population found on walnut trees. Although a perfect match between specific hybridization patterns and MLSA groups was not observed, it was clear that all the HPs corresponding to none (HP0) or a single positive *Xaj* specific markers (HP16, HP17, HP18) were correlated with more divergent MLSA groups composed by *X. arboricola* and *Xanthomonas* sp. isolates. Moreover, when MLSA and dot-blot hybridization patterns were combined, it was possible to enhance the genotyping resolution and disclose at least 31 divergent *Xanthomonas* lineages, including 17 lineages of *Xaj*, three lineages of *X. arboricola*, 11 lineages of *Xanthomonas* sp., among the 131 isolates associated with walnut trees.

Several important walnut production regions were included in this study, taking into consideration distinct bioclimatic conditions and different culture practices, namely Trás-os-Montes (e.g. Carrazeda de Ansiães location) with intensive summer drought and winter severity, i.e. mesomediterranean bioclimatic region, characterized by a walnut production largely based on isolated walnut trees; Alentejo (e.g. Beja and Estremoz locations) dominated by regularly warm temperatures and dry climate, i.e. supramediterranean bioclimatic region; and Minho (e.g. Ponte da Barca and Ponte de Lima) with high annual precipitation and relatively mild summers, i.e. mesotemperature bioclimatic region, both major walnut producing areas in Portugal characterized by orchards planted with selected *Juglans regia* cultivars. Although bioclimatic features and culture practices have been suggested to influence not only the epidemiology and prevalence of walnut disease, but also the bacterial population structure (9, 22, 23), our data show that some Xanthomonads lineages were not constrained by bioclimatic pressures and management strategies. In fact, *Xaj* lineages 1, 9, 12, and 14 appeared to be the most prevalent over time and uncover a widespread distribution in Portugal. On the other hand, some *Xaj* lineages were shown to have a narrower distribution, despite the exhaustive sampling effort (Lin6 and Lin17 specific to Alcobaça location; and Lin5 exclusively found in mesomediterranean bioclimatic region). These lineages could not be detected in other regions either because they are underrepresented in comparison with other prevalent *Xanthomonas* lineages, or because these clonal lineages are particularly adapted to specific ecological conditions. Future sampling efforts will be essential to track these lineages to determine their prevalence and to identify possible lineage specific adaptation traits. Moreover, seven lineages of *Xaj* (Lin2, Lin4, Lin8, Lin10, Lin13, Lin15, Lin16) and the majority of *X. arboricola* and *Xanthomonas <u>sp.</u>* genetic lineages described in this study (13/14 lineages – Lin18, Lin19, Lin20, Lin22, Lin23, Lin24, Lin25, Lin26, Lin27, Lin28, Lin29, Lin30, and Lin31) showed to be occasional, being the less frequent lineages.

Beyond the bioclimatic and geographic variables, host specific attributes namely organs of walnut trees, e.g. leaves, fruits, and trunks are expected to constitute distinct selective pressures which might favour the emergence of distinct *Xaj* lineages as previously reported (5, 24). Some studies hypothesized that genomic plasticity of *Xaj* confers a high adaptation to very different environmental niches namely through the gain of additional features which might have led to *Xaj* lineages as the one that have been associated with VOC disease (5). Furthermore, the same tree could be infected simultaneously by different *Xaj* strains, causing distinct symptoms (WBB, BAN and VOC) (5, 39). In the present study, we show that the diversity of bacterial xanthomonads found on walnut trees is more complex than originally thought, being characterized by distinct lineages of *Xaj, X. arboricola* and *Xanthomonas sp*. found to co-colonize the same walnut organ sample. Furthermore, we gathered evidence suggesting that co-colonization is not occasional, since was found in 32 of our 64 walnut trees sampled as shown by the coexistence of different *Xaj* lineages and *Xaj* lineages with *Xanthomonas* sp. lineages infecting leaves of the same walnut tree (Fig. 3). Interestingly, some *Xaj* lineages appeared strongly associated, as in the case of Lin1 frequently isolated with Lin12, and Lin6 recovered simultaneously with Lin11 in all leaf samples. Although, further investigations are important to determine if the xanthomonads diversity within the same walnut host plant and even within the same plant organ, is a mix of bacterial populations colonizing evenly the same host plant and organ, or if it results from the co-colonization of dominant versus lessened xanthomonads populations, it was recently proposed that sympatric populations, as pathogenic and nonpathogenic strains found together on walnut buds, may have important effects on genetic dynamics of new strains emergence (16, 20).

It is currently acknowledged that functional T3E have been pointed out as essential for pathogenicity in *X. arboricola* (20, 28, 29, 37). In fact, noninfective *X. arboricola* strains were characterized by the absence of genes encoding for some type III effectors proteins (T3Es) (15, 20, 28). When studying the presence of the T3Es genes *xopR, avrBs2, xopF1* and *xopN*, suitable to distinguish three groups of nonpathogenic strains (NP1, NP2, and NP3) according Essakhi et al. (20), we could assign most of the 19 isolates evaluated from cluster VII to X to nonpathogenic groups NP2 and NP3, with exception for two isolates (CPBF 78 and CPBF 424) which displayed a new pattern of T3E genes (presence of *xopR* and *xopF1*; absence of *avrbs2* and *xopN*) (Fig. 4). Moreover, when the pathogenicity of isolates with distinct composition of T3Es was assessed on leaves of walnut plantlets (CPBF 75, CPBF 367, CPBF 424, CPBF 1488, CPBF 1514), only strain CPBF 424, which was obtained from walnut buds, induced symptoms on their host of isolation. This result is particularly relevant and raise questions about the genomic virulence features of this *Xanthomonas* sp. strain (CPBF 424) on walnut trees and could be particular important for disclosing genome dynamics and for pathogenicity emergence of *X. arboricola*.

In conclusion, this study analysed the distribution of *Xaj* genetic diversity in Portugal, which consisted in extensive surveys conducted for three consecutive years, in distinct walnut producing regions characterized by diverse bioclimatic regions and different walnut production practices. Comprehensive genotyping analyses allowed to identify the most prevalent *Xaj* lineages, the possible emergence of new *Xaj* lineages and disclose non-infective *X. arboricola* strains and a non-*arboricola* pathogenic *Xanthomonas* sp., which might provide new insights to elucidate new *Xanthomonas* pathoadaptations.

## Materials and Methods

### Bacterial isolation from different walnut plant organs

A total of 94 walnut plant samples were collected from different plant organs (66 leaves, 17 fruits, four branches, six buds and one catkins) of 64 symptomatic walnut trees (*Juglans regia*) distributed throughout Portugal. Sampling were done in 14 geographic locations along four bioclimatic regions, characterized by distinct thermoclimatic parameters (32): mesotemperature - Mt (Ponte da Barca and Ponte de Lima locations), mesomediterranean - Mm (Carrazeda de Ansiães, Baião, Guarda and Seia locations), supramediterranean - Sm (Alcobaça, Beja, Bombarral, Estremoz, Leiria and Loures locations) and thermomediterranean - Tm (Azeitão and Portimão locations). Sampled walnuts were either isolated trees or walnut hosts found in orchards established with different French, American or/and Portuguese walnut cultivars, including F1 hybrids (Table 1). Sampling occurred between April and October (Leaves from June to October; Fruits from June to September; Branches in June and July; Catkins in September; Buds in April and September). Apart from one sample collected in 2009, all trees were sampled between 2014 and 2016, with the trees Jr#01, Jr#03, Jr#06, Jr#18, Jr#26 and Jr#27 being sampled in different years (Table 1). Necrotic lesions characteristic of WBB or BAN symptoms were observed in all leaves, fruits and branches sampled, and all buds and catkins samples collected were asymptomatic. Multiple samples of different organs were also collected at the same sampling date from the walnut trees Jr#02, Jr#03, Jr#05, Jr#07, Jr#08, Jr#11, Jr#18, Jr#25, Jr#26, Jr#27, Jr#30, Jr#35, Jr#47 and Jr#56 (Table 1).

Sample preparation for bacterial isolation was carried out differently for symptomatic and asymptomatic material: i) for symptomatic leaves, fruits and branches, plant tissues adjacent to necrotic areas were first excised using a sterile scalpel; ii) for asymptomatic buds and catkins, either single terminal buds, axillary bud groups or catkins groups of the same branch were excised also using a sterile scalpel according sampling procedures previously described (7, 8). Bacterial isolation was carried out as procedure detailed in Fernandes et al. (27). One to three isolates were selected per sample and stored at −80°C at the Portuguese Collection of Phytopathogenic Bacteria (CPBF - Colecção Portuguesa de Bactérias Fitopatogénicas, Oeiras, Portugal).

### Growth conditions of bacterial pure cultures and DNA extraction

The whole set of xanthomonads walnut isolates (Table 1) and *X. arboricola* strains used in this work were cultured at 28°C on YGC medium (5 g liter^−1^ yeast extract, 10 g liter^−1^ glucose, 30 g liter^−1^ CaCO_3_, 15 g liter^−1^ agar). Genomic DNA from pure bacterial cultures was extracted using the EZNA Bacterial DNA Purification kit (Omega Bio-Tek, Norcross, GA), following the manufacturer’s instructions, and quantified using the Qubit 2.0 Fluorometer HS Assay (Invitrogen, Carlsbad, CA).

### Multilocus sequence analysis and dot blot hybridization

Multilocus sequence analysis (MLSA) was carried out using the concatenated sequences of four housekeeping gene fragments: *acnB* (aconitase), *fyuA* (tonB-dependent receptor), *gyrB* (DNA gyrase subunit B) and *rpoD* (RNA polymerase sigma factor). Primer pairs used for *acnB* amplification (684 bp) were described by Parkinson et al. (34) and for *fyuA* (724 bp), *gyrB* (904 bp) and *rpoD* (915 bp) by Young et al. (35). The PCR reaction mixture (total volume of 40 µL) contained 1X DreamTaq Buffer with 2.0 mM MgCl_2_ (Fermentas, Ontario, Canada), 0.2 mM of each dNTP (Fermentas), 0.2 µM of each forward and reverse primers, 1U of DreamTaq DNA Polymerase (Fermentas) and 25 ng of bacterial DNA. PCR conditions for the four genes, were 95°C for 5 min, followed by 30 cycles of 95°C for 30 s, 54°C for 30 s, 72°C for 45 s, and a final extension step of 72°C for 10 min. PCR products were purified using the illustra GFX GEL Band Purification kit (GE Healthcare, Buckingham-shire, United Kingdom), according to the manufacturer’s instructions, and sequenced on both strands (STAB Vida, Caparica, Portugal). Consensus nucleotide sequences obtained for each gene fragments were aligned, trimmed and concatenated using the Geneious v. 9.1.7 software (Biomatters, Auckland, New Zealand). The 131 concatenated sequences (513 bp of *acnB*, 640 bp of *fyuA*, 828 bp of *gyrB* and 793 bp of *rpoD*) obtained from the Portuguese isolates were used to build a Maximum Likelihood tree based on the General Time Reversible (GTR+G+I) model in MEGA 7.0 (40). To account for the known *Xanthomonas* genomic diversity, 32 additional *X. arboricola* strains were included in the analysis together with 89 *Xanthomonas* spp. strains, for which all *acnB, fyuA, gyrB* and *rpoD* sequences were available in the GenBank database.

Dot blot assays were performed as described in a previous study using nine *Xaj* specific DNA markers (XAJ1 to XAJ9) (27). Briefly, 100 ng of bacterial DNA were bound to a nylon hybridization transfer membrane (Roche Diagnostics GmbH, Basel, Switzerland) using a Bio-Dot microfiltration unit (Bio-Rad, Hercules, CA). Hybridization was carried out overnight at 68°C to ensure high stringency, with each of the nine DIG-labelled probes (XAJ1 to XAJ9). Dot blot images were acquired using a Molecular Imager ChemiDoc system (Bio-Rad, Hercules, CA).

### Type three effector genes assessed by dot blot assays

The presence of four type three effector genes (T3E), *xopR, avrBs2, xopF1* and *xopN*, was assessed by dot blot hybridization assays. Genes *xopR, avrbs2, xopF1* have been considered to be ubiquitous T3E in strains of *X. arboricola*, whereas *xopN* has been suggested to be normally associated with *X. arboricola* strains from pathovars *juglandis, pruni* and *corylina* (15). Moreover, the distribution of these T3Es genes have been referenced to differentiate pathogenic from nonpathogenic strains of *X. arboricola* isolated from walnut trees (20). PCR primers used for preparation of T3E DNA probes were previously described (15). Partial sequences of the four T3E genes were obtained for *Xaj* strain LMG 751 using the PCR reaction conditions described above. PCR amplifications were performed with one cycle of 5 min at 95°C, followed by 35 cycles of 35 s at 95°C, 60 s at 60°C, 60 s at 72°C and a final step of 10 min at 72°C. Each DNA amplicon obtained (303 bp of *xopR*, 850 bp of *avrBs2*, 779 bp of *xopF1* and 864 bp of *xopN)* was purified with the illustra GFX GEL Band Purification kit, and sequenced (STAB Vida) to confirm its identity. The DIG-High Prime kit (Roche Diagnostics GmbH, Basel, Switzerland) was used for probe labelling, following the reference protocol available and a final probe concentration of 100 ng/ml was used in dot blot assays performed as described above. In addition to the 35 walnut xanthomonads isolates, one nonpathogenic strain of *X. arboricola* (CFBP 1022) and three *Xaj* reference strains (CFBP 176, LMG 747 and LMG 751) were also included in each dot blot assay.

### Pathogenicity assays

*Juglans regia* cv. Hartley seedlings were used for determination of pathogenicity of selected isolates. After 30 days of cold stratification treatment at 3-5°C to break dormancy, *Juglans regia* seeds were sown in sterilized sand substrate and germinated during 60 days at alternated temperatures, 16 hours day at 30°C and 8 hours night at 20°C (41). Walnut plantlets were then maintained in a climatic chamber under controlled environmental conditions of 16-hour photoperiod (16 h of light at 24°C and 8 h of darkness at 18°C).

Bacterial inoculations were performed when walnut plantlets had at least four young leaves fully expanded. Three plantlets were used for each isolate tested. Inoculum suspensions, prepared with sterile distilled water, were obtained from pure cultures grown on nutrient agar (NA) medium at 28 ± 2°C for 48 h. Bacterial suspensions were adjusted to a concentration of approximately 1 × 10^8^ CFU ml^−1^ and confirmation of bacterial inoculum concentration was carried out by plating serial decimal dilutions on NA medium, with viable cell counting made 48 h after incubation. Plantlets were inoculated by spraying with a manual atomizer until runoff and kept in closed polyethylene bags for 48 h to promote bacterial infection, under the same temperatures and photoperiod conditions mentioned above. Plastic bags were then opened and plants maintained during four weeks for development of symptoms. Walnut plantlets sprayed with sterile distilled water were used as negative controls. Positive controls were performed by spraying a suspension of the reference type strain *Xaj* LMG 747 and *Xaj* isolate CPBF 1480 using the same concentration of viable cells. In order to fulfil Koch’s postulates, reisolation was performed from leaves presenting necrotic spots (42).

### Accession number(s)

GenBank accession numbers corresponding to *acnB, fyuA, gyrB* and *rpoD* sequences of xanthomonads isolates from walnut is available as supplemental material (Table S1).

## Acknowledgments

This work was co-financed by the European Structural & Investment Funds (ESIFs) through the Operational Competitiveness and Internationalization Programme – COMPETE 2020 and by National Funds through FCT – Fundação para a Ciência e a Tecnologia, within the framework of the project EVOXANT (PTDC/BIA-EVF/3635/2014, POCI-01-0145-FEDER-016600). Camila Fernandes is supported by a fellowship from the FCT (SFRH/BD/95913/2013). The authors would like to thank the walnut producers who collaborated in this study, Lurdes Santos and Paulo Godinho for the work performed on walnut seedlings and Joaquim Trindade for taking the photos of pathogenicity results.

